## Supplementary figures and images for "Neuroprotective Effects of VEGF-B in a Murine Model of Aggressive Neuronal Loss With Childhood Onset"

### Supplemental Figure 1

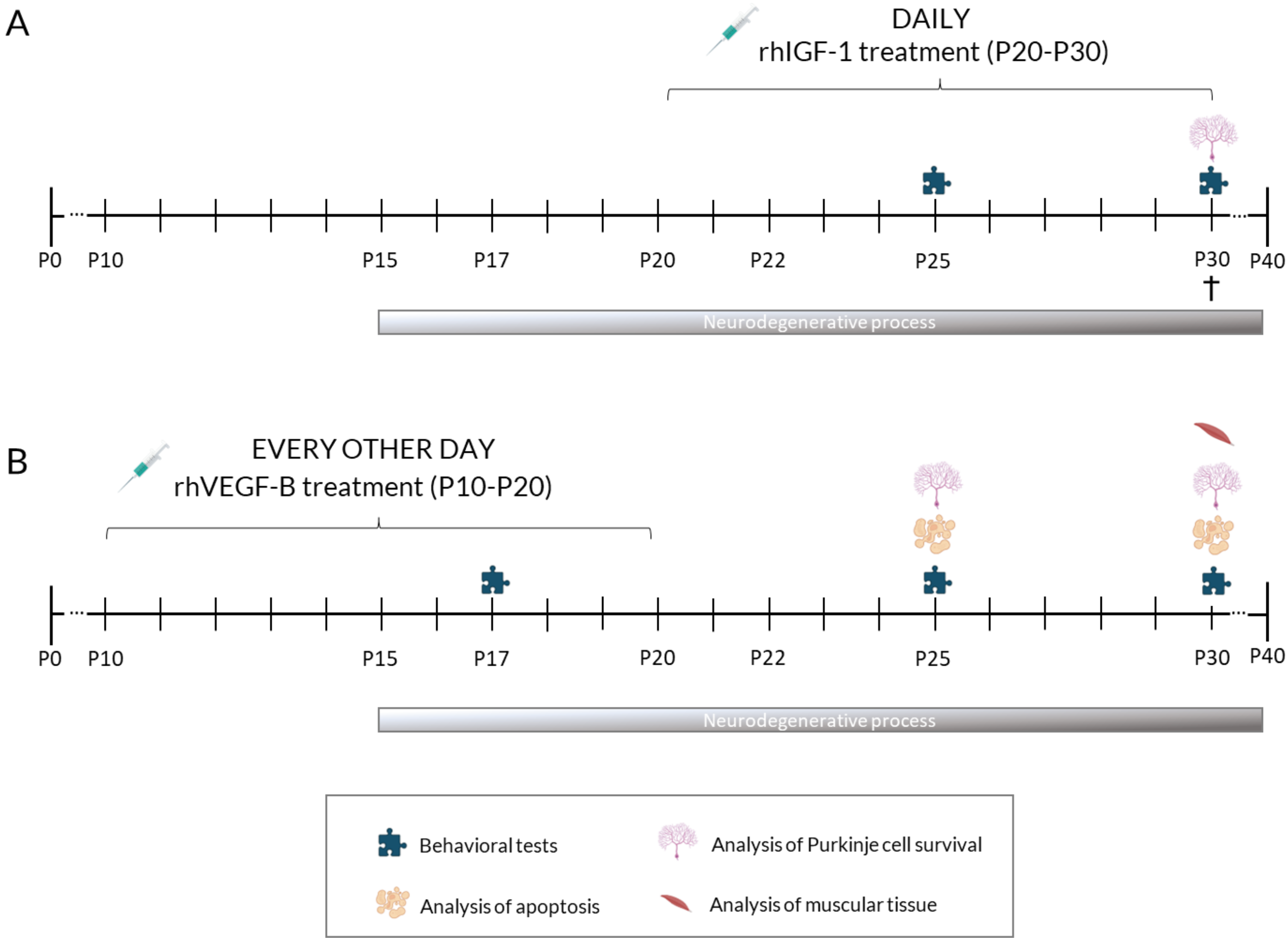
