## Supplemental Figure 1 for "Neuroprotective Effects of VEGF-B in a Murine Model of Aggressive Neuronal Loss With Childhood Onset"

**Figure S1**. **Experimental design of the different treatments evaluated in this study, the main analyses carried out and their relationship with the neurodegenerative process of the PCD mouse**. (A) Timeline details of rhIGF-1 administration, the time points for behavioral tests and tissue collection for Purkinje cell survival analyses. (B) Timeline details of rhVEGF-B administration, the timepoints for behavioral tests and tissue collection for Purkinje cell survival, apoptotic and muscular analyses.
